# Rapid and cost-effective production of high-quality insect genome sequences

**DOI:** 10.1101/2024.10.06.616902

**Authors:** Joshua Gilligan, Sarah N. Inwood, Tom W.R. Harrop, Phoebe Keddell, Kate McPhail, Connor J. Sproston, Darren W. Williams, Gertje Petersen, Anton Hovius, Julia Kasper, Joseph Guhlin, Peter K. Dearden

## Abstract

Insects are the most diverse group of animals on earth, with nearly 1 million species described to date, yet this variation is not reflected within the ranks of genomes currently available. In the current age of genetics, the generation of high-quality reference genomes is invaluable, often necessary and expensive. We prove that insect genomes do not need to come at a large cost. We demonstrate a cost-efficient method for producing high-quality insect genomes, with comparisons to current reference assemblies.

## Introduction

Insects are the most diverse group of animals on earth, with nearly 1 million species described to date^1^ and predictions placing the total number of species at 6-12 million^2,3^. This abundance of genera is reflected in the diversity of size, ploidy and complexity of insect genomes. Out of the 11,118 metazoan genomes currently found in GenBank, however, only 3,651 are insects (as of 13/Apr/2023). Genomes of a broad range of insect species, is crucial to understanding the breadth of their variation, however developing genomes is expensive and time-consuming.

High-quality genomes form the basis of all areas of genomic research. They are needed to ensure that most of the within-species genomic variation is represented in published reference sequences, vital for genome diversity studies within and between species. For insect breeding or conservation studies, high-quality references are needed to map genetic variation associated with important traits^4^. With regulatory evolution thought to be responsible for many micro-evolutionary changes between species^5^, high-quality genomes can also enable the identification of the whole gene complement and potential regulatory elements. While research into individual species is set to benefit greatly from the availability of good quality genomes, a greater **number of** genomes from different orders would help investigations of the evolution of life across nature’s most **speciose** animal group.

Here we present a simple, rapid method for the production of insect genomes that is relatively cheap, and yet produces high-quality results with few gaps. We use this technique to sequence and assemble the genomes of 8 species, demonstrating both the advantages and limitations of this approach.

## Methods

### Samples

*Apis mellifera* Linnaeus, 1758 (Honeybee in the following) (Hymenoptera) samples were provided by Taylor’s Pass Honey Company from Apiaries in Marlborough, New Zealand. *Polistes dominula* (Christ, 1791) (European paper wasp) (Hymenoptera) samples were collected by the authors in Galloway, Alexandra, Central Otago, New Zealand (45.20901° S, 169.46064° E.). *Culex quinquefasciatus* Say, 1823 (Southern house mosquito (Diptera) was collected from Napier by Ian Jarvis, *Anatalanta aptera* Eaton, 1875 (Diptera) was provided by David Renault. *Boreoides tasmaniensis* Bezzi, 1922 (Tasmanian wingless soldier fly) (Diptera) was collected by authors in Dunedin, New Zealand (45.87623° S, 170.48032° E.) *Teleogryllus commodus* (Walker, 1869) (Black field cricket) (Orthoptera) were purchased from InZect direct (https://inzectdirect.co.nz) and *Celatoblatta quinquemaculata* (Johns, 1966) (Otago alpine cockroach (Blattodea) was collected from the Rock and Pillar Range, Otago (45.53488° S, 169.94185° E) by Dr Craig Marshall.

All insects were frozen whole and stored at -20 or -80 degrees centigrade before DNA extraction.

### DNA extraction

A single whole insect, or in the case of honeybees, an insect head, was ground in liquid nitrogen in a mortar and pestle. The substrate was then transferred to 2 ml of 800 mM guanidine HCl, 30 mM Tris-CL pH 8.0, 30 mM EDTA, 5% Tween-20, 0.5% Triton X-100, containing 200 mg/ml RNase A and 1 mg/ml Proteinase K. The buffer-substrate mixture was mixed by inversion and incubated at 50 degrees for at least two hours with occasional additional inversion.

After digestion, the liquid was transferred to a Qiagen Genomic-tip 20 (Qiagen, Venlo) and DNA was extracted as per the manufacturer’s protocols.

DNA concentration was determined using a Qubit (Thermo Fisher, Waltham, MA) high-sensitivity kit.

### RNA extraction

RNA was extracted from dissected cricket oocytes using a Trizol-RNAeasy protocol previously described for extraction from bee ovaries^6^. Purified RNA was quantified using a Qubit RNA kit (Thermo Fisher, Waltham, MA).

### DNA sequencing library construction

Where DNA concentrations were too low (less than 1 μg in total) for library production, the DNA was amplified using the Qiagen Repli-G MIDI whole genome amplification kit (Qiagen, Venlo) following protocols provided by Oxford Nanopore technologies. Herein, 10 pg -1 ng of genomic DNA in 5 μl volume were incubated briefly with 5 μl DLB buffer from the Repli-G MIDI kit for 3 minutes at room temperature, before 10 μl of Repli-G MIDI stop buffer were added. This mixture was transferred to a 0.2 ml PCR tube and a mixture of 29 μl Repli-G MIDI reaction buffer and 1 μl REPLI-g MIDI DNA Polymerase was added. Each sample was incubated for 16 hours at 30° C before the enzyme was denatured at 65° C for 3 minutes. The resulting DNA was purified using AMP pure XP magnetic beads (Beckman Coulter, Brea, CA), and then quantified using the Qubit high-sensitivity kit.

A total of 1.5 μg of amplified DNA was treated with 15 U of T7 Endonuclease I in 5 mM NaCl, 1 mM Tris-HCl, 1 mM MgCl2, 0.1 mM DTT, pH 7.9 for 15 minutes at 37°C. Debranched DNA was purified using AMP pure XP magnetic beads in 0.1 M Tris-HCl, 1 mM EDTA, 1.6 M NaCl, and 11% PEG 8000. DNA was quantified using the Qubit high-sensitivity kit and 700 ng – 1 μg were used in subsequent DNA preparation.

Non-amplified genomic DNA was sheared before library preparation, as we have found that tends to increase overall flow cell lifetime, except for that from honeybees where this was found to be unnecessary. Shearing was performed using Covaris g-TUBE (Covaris, Bankstown, NSW) following the manufacturer’s instructions.

Oxford Nanopore library preparation was carried out using either the SQK-LSK109 (honeybee and *Polistes dominula*) or SQK-LSK110 ligation sequencing kits (all other species) following the manufacturer’s instructions (Oxford Nanopore Technologies, Oxford). For non-amplified samples, DNA added to the ligation sequencing reaction ranged from 0.2 – 1 μg depending on what was available.

### DNA sequencing

Libraries were quantified and loaded onto 9.4.1 FLOMIN-106 Oxford Nanopore flow cells following the manufacturer’s instructions. These were then run using MinKNOW (most up-to-date versions used at the time of sequencing) software, without real time base-calling, aiming for at least 8 GB of sequence for most species. For honeybees, this was easily achieved and three drone samples could be run on one flow cell. For most other species, two flow cells were used. When both amplified and un-amplified samples were being sequenced, one flow cell was used for amplified, and one for unamplified. Flow cells were stopped and cleaned when active pore numbers dropped below 50, using the Oxford Nanopore Flow Cell Wash Kit, reloaded with new aliquots of sequencing libraries and restarted. Flow cells and libraries were used until exhausted to get as much data as possible for each species.

### RNA sequencing

RNA sequencing libraries were produced using the Illumina Truseq Stranded mRNA process and sequenced on an Illumina HiSeq 2500 (Illumina, San Diego, CA) by the Otago Genomics Facility. BBDuk v39.01^8^ was used to quality trim RNA-seq reads and to remove Illumina sequencing adapters, using default settings with trimq set to 15. FastQC 0.11.5^7^ was then used to verify adapter removal and the quality of trimmed reads. Trimmed reads from all samples were then concatenated, and STAR v2.7.11b^9^ was used to map reads using a two-pass approach.

### Base-calling and assembly

*Culex* sample was base-called earlier using Guppy(6.0.1+652ffd1) using the sup accuracy base calling model, and was trimmed for adaptors using Porechop(0.2.4)^10^

Fast5 files were base-called using Guppy (6.4.6) using the sup accuracy base calling models. Base-called data was trimmed for adaptors using Porechop_abi (0.5.0)^10^

Genome assembly was performed using Flye (2.9.2-b1786)^11^, with both the nanoraw and nanohq setting. Assembly polishing was carried out using Medaka (1.8.1), using the pore chopped reads as data input.

### Data visualisation and analysis

Genome completeness was assessed using BUSCO (5.4.7)^12,13^. Genome assemblies were visualised using BlobToolKit^14^ and Genome comparisons were carried out using D-GENIES (1.2.0)^15^.

Read depth was estimated using mosdepth (0.2.6)^16^.

## Results and Discussion

### Re-assembling honeybee genome

To test our assembly techniques, sequencing of honeybee genomes was performed initially, since *Apis mellifera* has a high-quality reference sequence available. First published in 2006^17^ (based on Sanger sequencing data), the genome of the honeybee has been extensively re-sequenced and revised as technology has advanced. The latest genome is AmelHAv3.1, which used PacBio; 10X Chromium; Phase Genomics HiC and BioNano sequencing and was then assembled with FALCON v. 0.5.0; Arcs v. 1.0.1; Links v. 1.8.5 and BioNano Solve v. 3.1, resulting in 192x genome coverage^18^.

Honeybees display haplodiploidy where males are haploid, while females are diploid. To provide the ideal conditions for our assembly technique we used single haploid male bees (drone) for each extraction, sequencing and assembly, sequencing three drones to determine the accuracy of our assemblies (compared to the reference) and variation produced by using different starting drones.

All three drones used in this analysis came from separate hives from a commercial bee operation and are likely to be related samples. To identify the most effective methods for de-novo assembly of this data we tested the nanohq and nanoraw settings of Flye^11^, with and without polishing with Medaka. Assembly statistics from various de-novo assembly options of this drone sequencing data are shown in Table 1 compared to the AmelHAv3.1 reference genome. All three genomes showed broadly equivalent assembly statistics to AmelHAv3.1, and the different assembly methods used make little difference to the completeness of the assembly.

**Table 1:** Assembly statistics from drone *Apis mellifera* de-novo assemblies.

| Name | Sum | N contigs | Average | Largest | N50 | Gaps |
| --- | --- | --- | --- | --- | --- | --- |
| TP1 hq | 231,416,688 | 448 | 516,555.1 | 13,878,957 | 7,940,090 | 20 |
| TP1 hq medaka | 231,691,222 | 448 | 517,167.9 | 13,895,631 | 7,949,490 | 0 |
| TP1 raw | 230,161,495 | 474 | 485,572.8 | 11,550,891 | 5,291,263 | 11 |
| TP1 raw medaka | 230,433,417 | 474 | 486,146.5 | 11,566,051 | 5,298,811 | 0 |
| TP2 hq | 229,193,540 | 604 | 379,459.5 | 11,629,068 | 5,112,234 | 24 |
| TP2 hq medaka | 229,479,622 | 604 | 379,933.2 | 11,643,953 | 5,118,903 | 0 |
| TP2 raw | 227,497,337 | 668 | 340,564.9 | 9,958,579 | 4,012,892 | 14 |
| TP2 raw medaka | 227,774,286 | 668 | 340,979.5 | 9,971,719 | 4,017,854 | 0 |
| TP3 hq | 228,117,322 | 487 | 468,413.4 | 13,575,959 | 7,257,561 | 20 |
| TP3 hq medaka | 228,388,983 | 487 | 468,971.2 | 13,594,565 | 7,267,892 | 0 |
| TP3 raw | 227,146,150 | 513 | 442,780 | 11,446,847 | 5,068,895 | 11 |
| TP3 raw medaka | 227,430,850 | 513 | 443,335 | 11,462,710 | 5,076,632 | 0 |
| **GCA_003254395.2_Amel_HAv3.1** | **225,251,012** | **177** | **1,272,605** | **27,754,200** | **13,619,445** | **51** |

To test the completeness of these assemblies, we carried out BUSCO^12,13^ analysis (Supplemental Table 2) comparing the BUSCO scores with the reference AmelHAv3.1 using snail plots to visualise the data (Figure 1 for TP1 assemblies, all others in Supplemental Figure 1). While all three genomes showed slightly worse BUSCO scores than the reference, these differences are minor and considering the lack of short reads, reasonable and expected.

**Figure 1:**
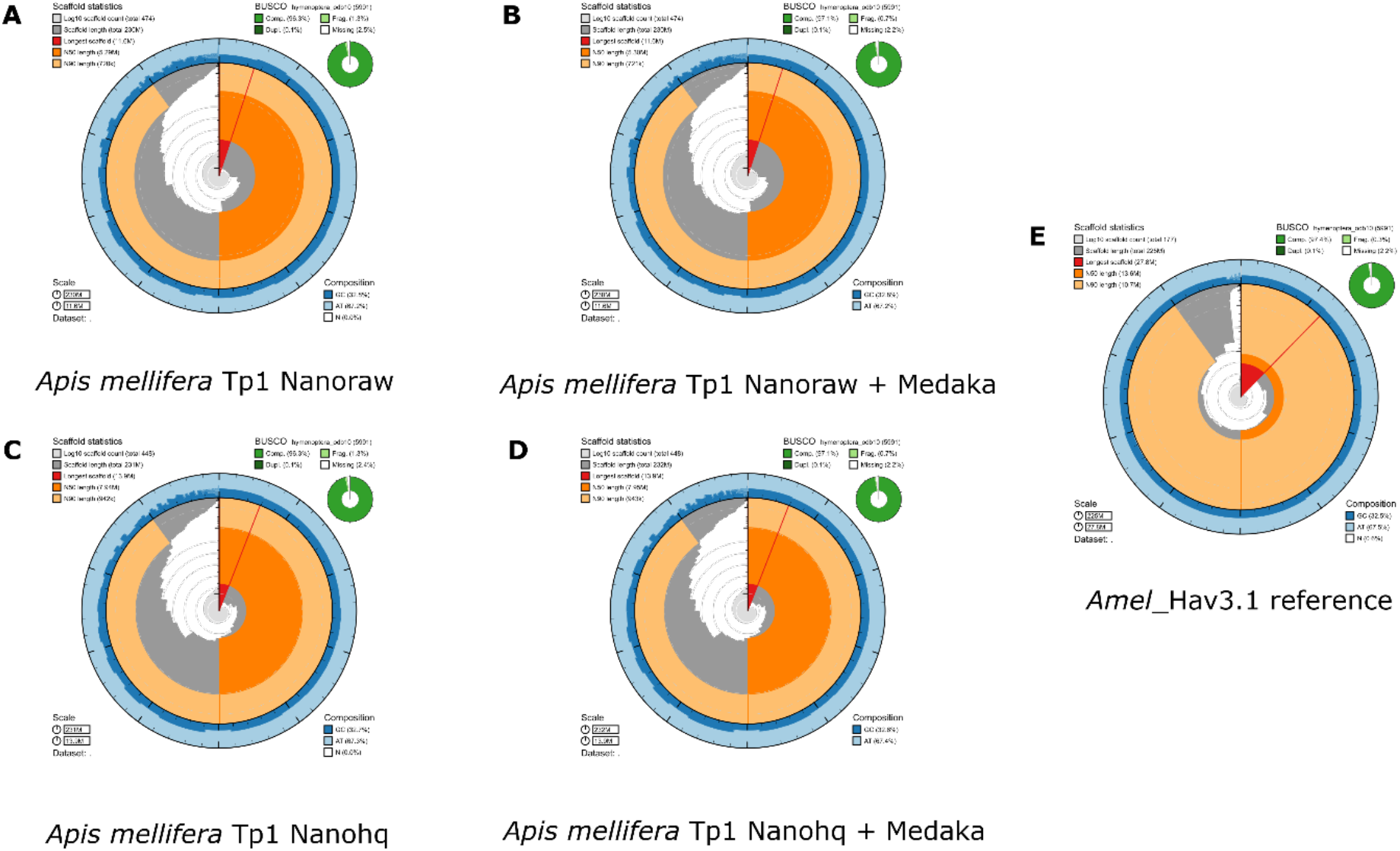
BlobToolKit Snail plot of Honeybee genomes. These plots show assembly metrics and BUSCO results. The main circumference of the plot is divided into 1000 size-ordered bins representing 1% of each genome. Scaffold length distributions are shown in dark grey, with the radius scaled to the longest scaffold (shown in red). Orange shows N50 scaffold length, tan shows N90 scaffold length. The light grey spiral shows the scaffold count on a log scale, with white bars indicating orders of magnitude. The dark/light blue ring indicates GC richness. BUSCO data is displayed at the right-hand corner, indicating the dataset tested against. A) Shows the honeybee drone TP1 assembled with nanoraw settings, B) TP1 with nanoraw and medaka polishing, C) TP1 with nanohq settings, and D) TP1 with nanohq and medaka polishing. E) shows the Amel_Hav3.1 reference genome

To determine the relationship between our various assemblies and the reference we carried out D-GENIES^15^ analyses to compare these genomes. Comparisons between different assembly approaches are shown in Supplemental Figure 2, and a comparison of one of the newly produced honeybee genomes (TP1 nano raw medaka) with AmelHAv3.1 via D-GENIES^15^ is shown in Figure 2. Figure 2 indicates that 96.83% of the genome aligns with the reference, a reasonable number considering the reference is derived from the DH4 line of bees from the USA and will not perfectly match honeybees from NZ.

**Figure 2:**
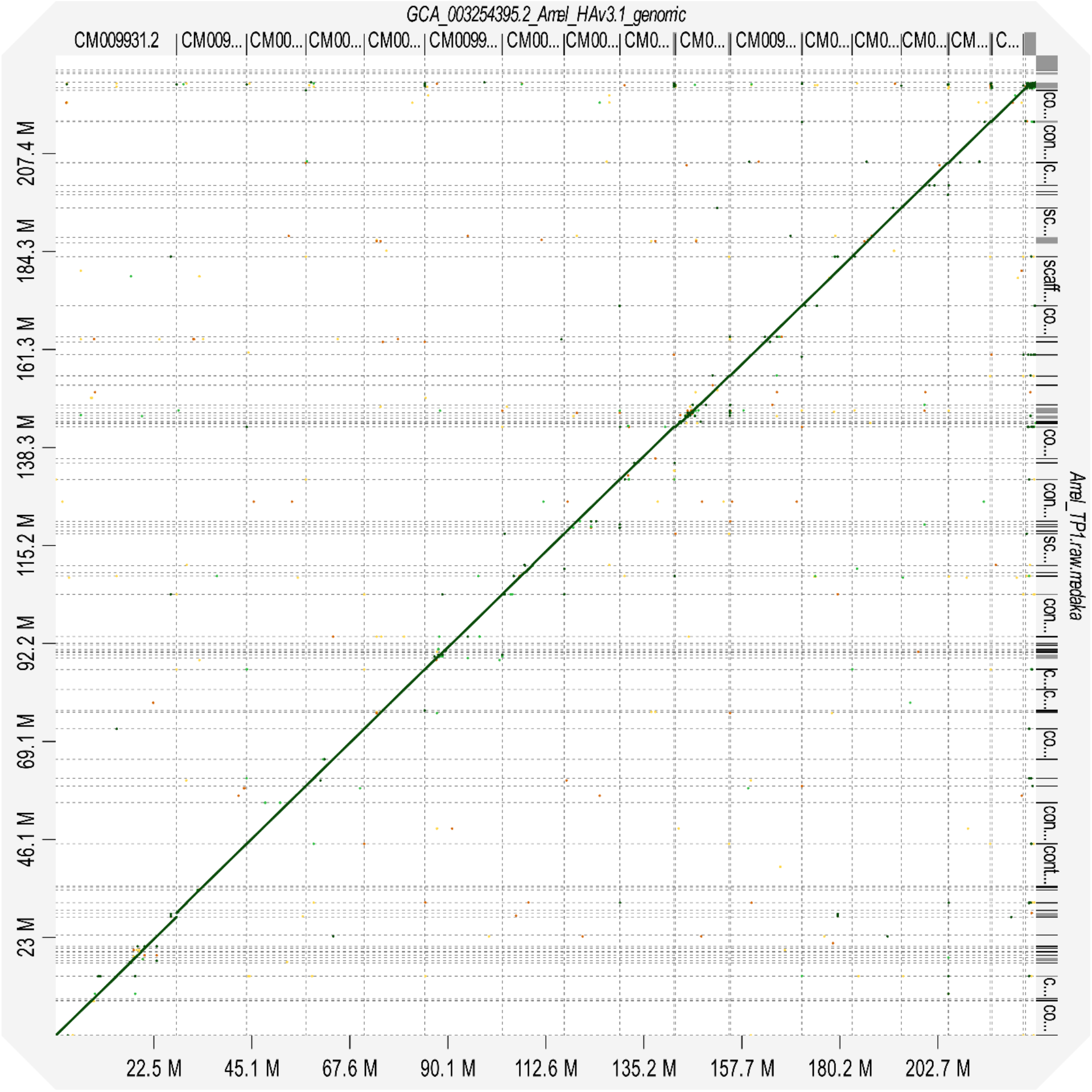
D-GENIES dot plot showing whole genome alignment of honeybee drone TP1 nanoraw medaka polished assembly (Y axis) against the Amel_Have3.1 reference genome (X axis). Ticks on each axis indicate contigs. The green line along the diagonal axis indicates strong alignment between the two assemblies.

We have generated a rapid and relatively inexpensive method for producing high-quality genomes for haploid honeybee drones. From start to finish, this work can be completed in a week (depending on computing resources) and generates genomes with BUSCO scores comparable to the intensively sequenced reference genome and with few differences in structure. The low-cost generation of high-quality genome sequence data from drones supports honeybee breeding programmes either in the reconstruction of queen genotypes^19^ or as reference genomes to support genotyping and diversity assessment^4,20^.

### Generation of genomes for other haplodiploid species

Given the high-quality honeybee genomes produced by our relatively inexpensive methods, this technique was applied to other insect species with haploid males. We focused on two species of invasive wasp in New Zealand, *Polistes dominula*, for which a previous genome is available^21^, and *P. chinensis*.

Assembly stats for these genomes, compared to the current *P. dominula* genome, are shown in Table 2. Using the Flye nanohq settings for assembly of the *P. chinensis* genome generated a genome size double that expected and much poorer quality than the genome generated by the nanoraw settings.

**Table 2:** Assembly statistics from *Polistes* de-novo assemblies.

| Name | Sum | N contigs | Average | Largest | N50 | Gaps |
| --- | --- | --- | --- | --- | --- | --- |
| Pchi hq | 380,215,862 | 5076 | 74,904.62 | 11,324,464 | 2,934,900 | 89 |
| Pchi hq medaka | 379,929,108 | 5076 | 74,848.13 | 11,344,732 | 2,937,056 | 3 |
| Pchi raw | 219,816,972 | 499 | 440,515 | 10,405,675 | 5,509,776 | 5 |
| Pchi raw medaka | 220,163,884 | 499 | 441,210.2 | 10,422,591 | 5,521,215 | 0 |
| Pdom hq | 235,319,780 | 463 | 508,250.1 | 11,430,933 | 5,954,638 | 8 |
| Pdom hq medaka | 235,653,755 | 463 | 508,971.4 | 11,441,828 | 5,967,432 | 1 |
| Pdom raw | 221,938,340 | 384 | 577,964.4 | 10,450,949 | 5,929,758 | 2 |
| Pdom raw medaka | 222,299,396 | 384 | 578,904.7 | 10,466,676 | 5,940,336 | 0 |
| **GCF_001465965.1_Pdom_r1.2_genomic** | **208,026,220** | **1483** | **140,273.9** | **7,126,315** | **1,625,592** | **14286** |

BUSCO scores and other data for these genomes (polished and unpolished) are shown in Supplemental Table 2, Figure 3, and Supplemental Figure 2.

**Figure 3:**
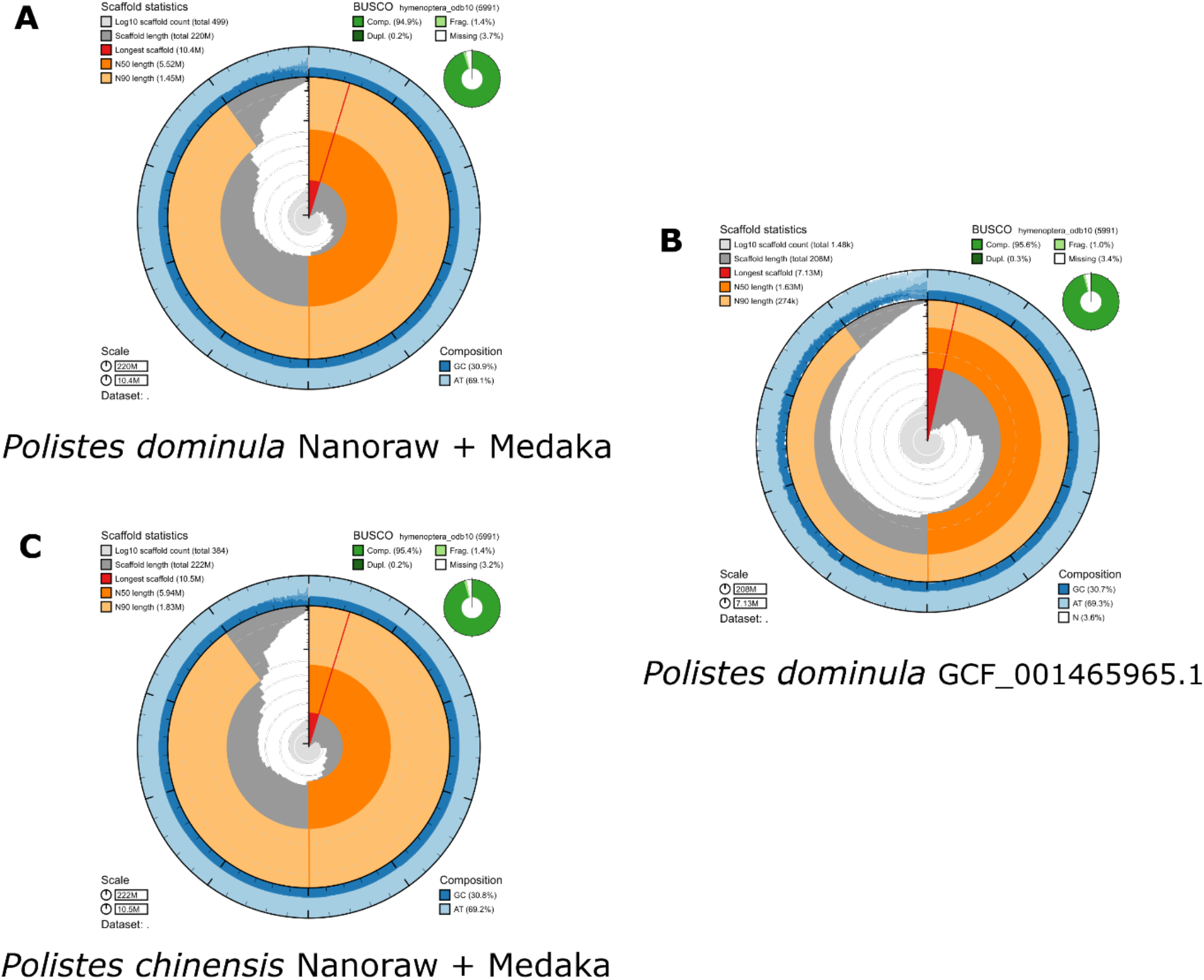
BlobToolKit snail plot of *Polistes* genomes. A) an assembly of *Polistes dominula* produced with nanoraw settings and medaka polishing. B) the current *P. dominula* reference. C) an assembly of *P. chinensis* produced with nanoraw settings and medaka polishing.

Figure 4 shows D-GENIES alignment of our *P. dominula* genome against the reference *P. dominula* genome (NCBI accession GCF_001465965.1). The agreement between these genomes, built using different techniques and data sources, is encouraging that our approach is effective. In addition, our best *P. dominula* genome has better assembly statistics and the same BUSCO score as the reference.

**Figure 4:**
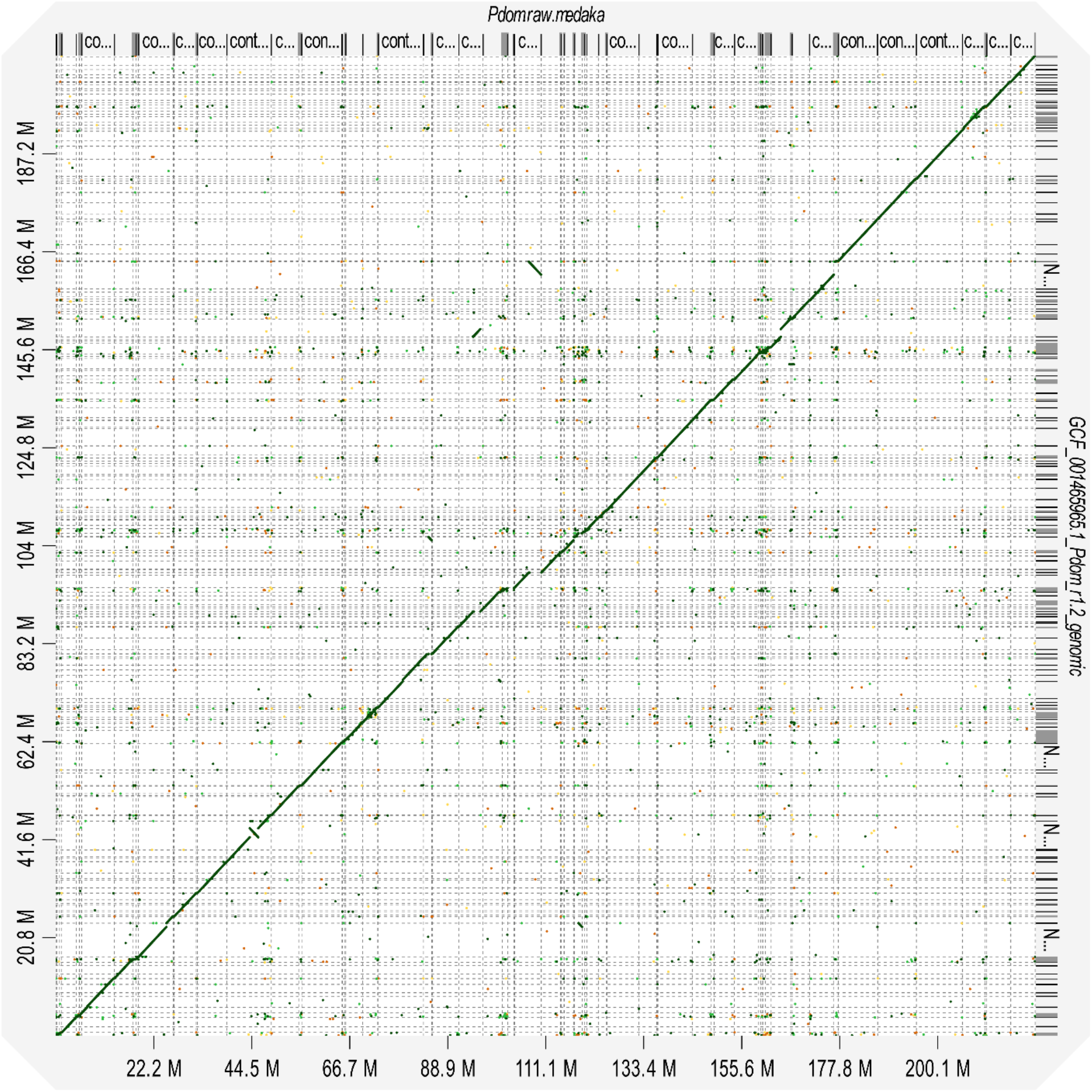
D-GENIES dot plot showing whole genome alignment of a nanoraw medaka polished *Polistes dominula* assembly (X axis) against the *P. dominula* GCF_001465965.1 reference assembly (Y axis). Ticks on each axis indicate contigs.

The different settings in Flye^11^, nanoraw vs nanohq, produced very different genomes for *P. chinensis*. Because of this, and the speed at which these assembly techniques can be applied, we would advise that both approaches be tested on novel genomes, using BUSCO^13^ and genome alignment methods as a metric for genome completeness and correctness.

These data indicate that the method of genome assembly presented here is effective in insect species beyond honeybees, can produce genomes of the quality or better than those currently used as references, and can produce plausible genomes from species currently without a reference.

This low-depth approach for de novo genomes highlights the practicality of sequencing less per individual, and instead sequencing multiple individuals for the same relative cost. This efficiency allows haplodiploid pangenomes to be generated more easily than ever before and could rapidly expand our ability to probe the relationship between structural variation and evolution.

### Generating genomes of diploid species

Given the success of building insect genomes using haploid individuals, we next attempted, using the same technology, to sequence the genomes of several diploid insects. These included *Culex quinquefasciatus* (a cosmopolite bloodsucking fly), for which a reference genome is available for comparison^22^, as well as several insects currently without published reference genomes: *Anatalanta aptera* (a wingless fly on dung), *Boreoides tasmaniensis* (a nectar feeding fly with wingless females), *Teleogryllus commodus* (a cricket and pasture pest in Australasia) and *Celatoblatta quinquemaculata* (a high-altitude New Zealand native cockroach). These species were chosen as representative examples of small and large diploid insect genomes, but also for their relevance in food production systems, since high-quality genomes would support further research.

Genome assembly statistics for all species are shown in Table 3 and BUSCO statistics for each genome in Figure 5 and Supplementary table 3. Some species (*B. tasmaniensis, A. aptera* and *T. commodus*), showed good results (especially BUSCO), but *Culex quinquefasciatus* and *C. quinquemaculata* showed low BUSCOs, low N50 and short scaffolds. For genomes that are well resolved (*B. tasmaniensis, A. aptera* and *T. commodus*) we consistently get between 1 and 4% missing BUSCOs, implying that these genomes are relatively complete. Since no reference genome is available for *T. commodus*, independently generated RNAseq reads were mapped directly to the assembled genome using STAR v2.7.11b^9^ in order to provide some information on the completeness of the genome.

**Table 3.** Assembly statistics for diploid genomes.

| Name | Sum | N Contigs | Average | Largest | N50 | Gaps |
| --- | --- | --- | --- | --- | --- | --- |
| *Antalanta* hq fasta | 825,923,543 | 38,981 | 21,187.85 | 4,413,887 | 140,503 | 298 |
| *Antalanta* hq medaka fasta | 828,625,753 | 38,981 | 21,257.17 | 4,414,452 | 141,667 | 5 |
| *Antalanta* raw fasta | 669,788,387 | 22,579 | 29,664.22 | 3,785,075 | 212,934 | 202 |
| *Antalanta* raw medaka fasta | 673,075,915 | 22,579 | 29,809.82 | 3,785,716 | 214,754 | 5 |
| *Boreoides* hq fasta | 587,859,249 | 3,050 | 192,740.74 | 5,659,661 | 816,210 | 99 |
| *Boreoides* hq medaka fasta | 588,337,548 | 3,050 | 192,897.56 | 5,660,189 | 816,930 | 4 |
| *Boreoides* raw fasta | 575,573,640 | 3,613 | 159,306.29 | 4,078,180 | 688,863 | 75 |
| *Boreoides* raw medaka fasta | 576,116,054 | 3,613 | 159,456.42 | 4,082,043 | 690,233 | 3 |
| *Celatoblatta* hq fasta | 3,390,745,100 | 122,180 | 27,752.05 | 392,267 | 39,786 | 281 |
| *Celatoblatta* hq medaka fasta | 3,425,007,452 | 122,180 | 28,032.47 | 393,179 | 40,232 | 71 |
| *Celatoblatta* raw fasta | 3,334,152,136 | 141,846 | 23,505.44 | 230,668 | 32,422 | 228 |
| *Celatoblatta* raw medaka fasta | 3,372,260,554 | 141,846 | 23,774.1 | 231,229 | 32,779 | 51 |
| *Culex* hq fasta | 863,248,721 | 21,916 | 39,388.97 | 442,262 | 62,047 | 143 |
| *Culex* hq medaka fasta | 866,861,681 | 21,916 | 39,553.83 | 442,922 | 62,187 | 24 |
| *Culex* raw fasta | 831,921,800 | 24,423 | 34,063.05 | 465,412 | 53,379 | 158 |
| *Culex* raw medaka fasta | 835,892,533 | 24,423 | 34,225.63 | 466,909 | 53,591 | 44 |
| *Teleogryllus* hq fasta | 2,143,040,480 | 26,954 | 79,507.33 | 5,327,702 | 400,140 | 125 |
| *Teleogryllus* hq medaka fasta | 2,153,693,185 | 26,954 | 79,902.54 | 5,333,068 | 401,290 | 7 |
| *Teleogryllus* raw fasta | 2,004,124,109 | 23,439 | 85,503.82 | 2,731,734 | 353,213 | 110 |
| *Teleogryllus* raw medaka fasta | 2,014,919,744 | 23,439 | 85,964.41 | 2,733,435 | 354,055 | 5 |

**Figure 5:**
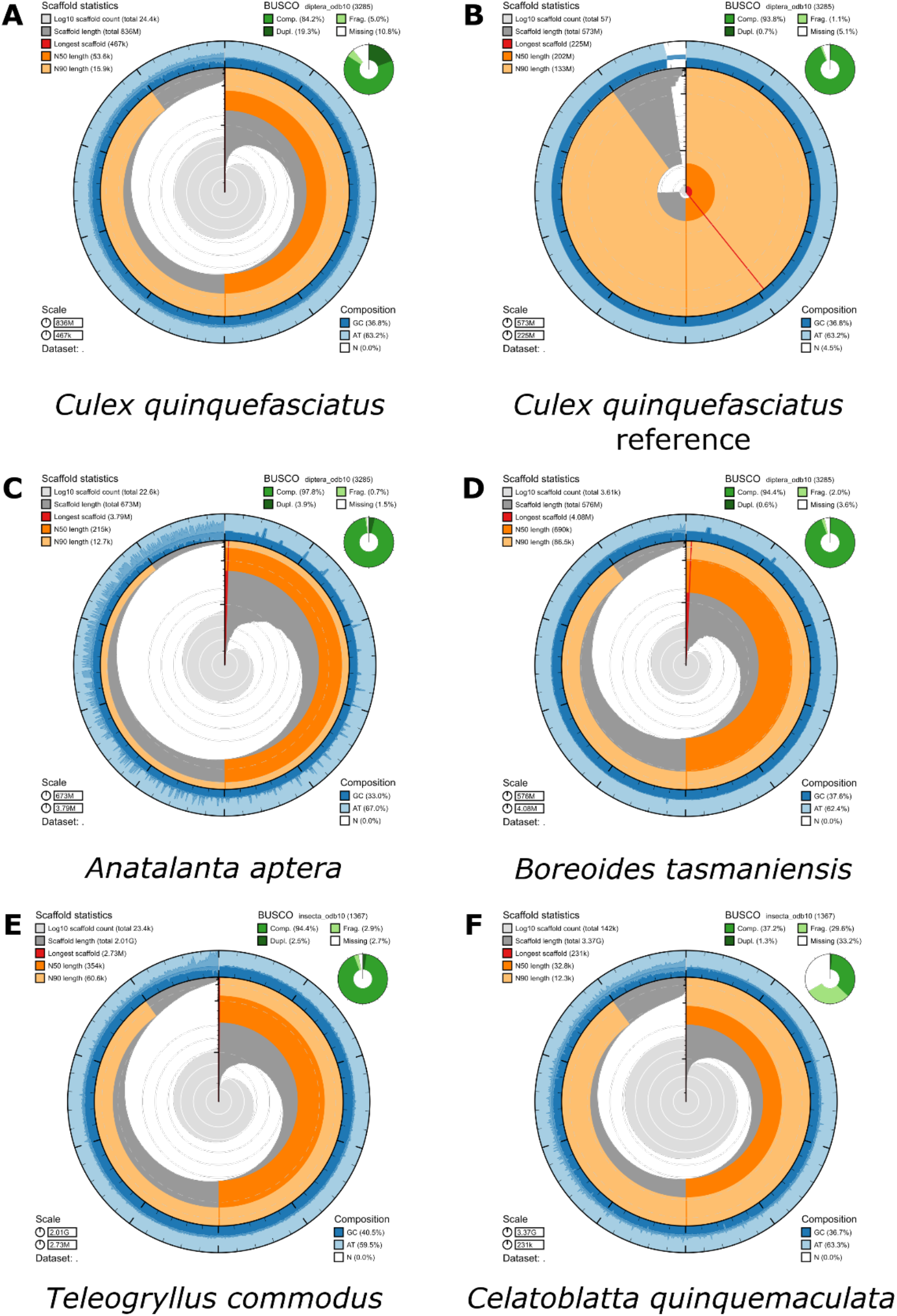
BlobToolKit snail plot of Diploid assemblies. A) an assembly of *Culex quinquefasciatus* produced with nanoraw settings and medaka polishing. B) the current *C*.*quinquefasciatus* GCF_015732765.1 reference. The genome we have generated has much worse assembly statistics and BUSCO scores. C) an assembly of *Anatalanta aptera* produced with nanoraw settings and medaka polishing. D) an assembly of *Boreoides tasmaniensis* produced with nanoraw settings and medaka polishing. E) an assembly of *Teleogryllus commodus* produced with nanoraw settings and medaka polishing. F) an assembly of *Celatoblatta quinquemaculata* produced with nanoraw settings and medaka polishing.

Mapping *T. commodus* RNAseq reads against the assembled genome resulted in a 93.8% read mapping (82.5% uniquely mapped, 11.3% mapped to multiple loci), consistent with results we get for mapping honeybee reads against the honeybee reference genome.

The *T. commodus* genome generated here is large (∼2 GB) and GC rich (40.5%), indicating that the method presented here can deal with challenging genome sequences.

### Failed genomes

For two of the species tested, we failed to produce sensible assemblies. For *Culex quinquefasciatus*, the numbers of missing BUSCOs were in the 9-10 % range, but duplicated BUSCOs reached 22.7%, whereas for *Celatoblatta quinquemaculata* missing BUSCOs ranged from 32%-34% depending on the assembly method. Alignment of our *Culex* genome with the reference genome (NCBI GCF_015732765.1) indicates our genome, while showing some resemblance to the reference, is fragmented and misassembled (Figure 6).

**Figure 6:**
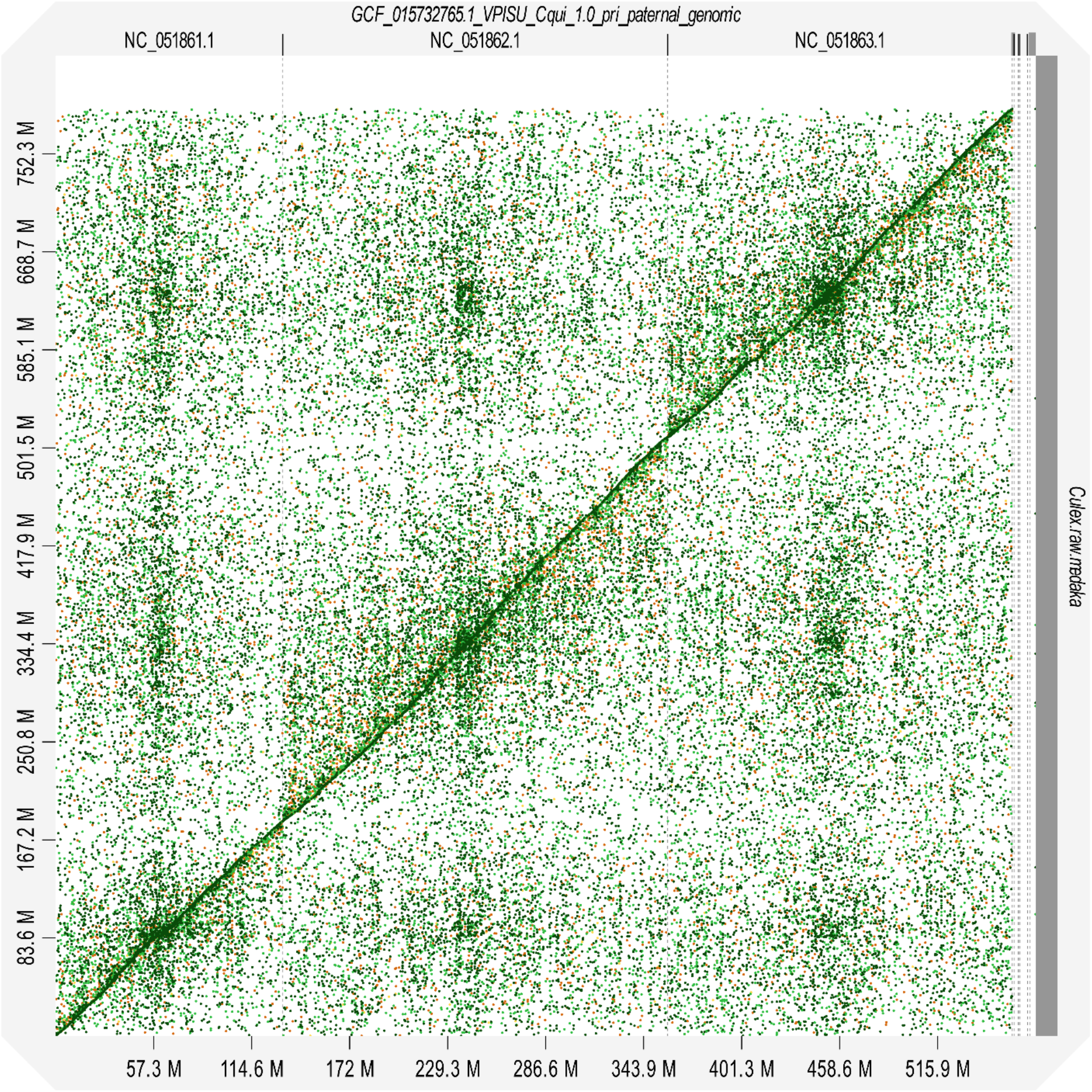
D-GENIES dot plot showing whole genome alignment of a nanoraw medaka polished *Culex quinquefasciatus* assembly (Y axis) against the *C. quinquefasciatus* GCF_015732765.1 reference assembly (X axis). Ticks on each axis indicate contigs.

Examining the coverage generated for our diploid genomes, it is not clear why these poor assemblies are occurring. It seems likely that this is not due to low coverage of the genomes. Coverage of *Antalanta aptera* and our *Polistes* assembly is between 18 to 50 fold, while for *C. quinquefasciatus* we have 10 fold and the *C. quinquemaculata* an estimated 8 fold coverage. Our *Teleogryllus commodus* genome however has an estimated 7 fold coverage, but generated a good assembly based on BUSCO and RNA-seq mapping results.

## Conclusions

Here we present a method that produces good-quality genomes for insects from Oxford Nanopore sequencing data alone. This method can assemble haploid and diploid genomes, and copes with large sizes and high GC content. The quality of our genomes seems to be reliant on coverage but intrinsic factors to the genome can impact our assemblies.

During the writing process, new base-calling algorithms were developed by ONT, with each of these iterations we were able to rebase call our original fast5 reads and make better genomes each time. With further advancements in Nanopore technology (kit 14 chemistry, new base calling algorithms and duplex stereo base calling), our ability to generate *de novo* genomes will only improve, rapidly cataloguing the diversity of arthropods with ease.

Our data also shows that polishing using Medaka can improve the contiguity of these genomes but makes little difference to BUSCO scores. Genome polishing would be important for genome alignments and exploring synteny, but is probably not necessary in the search for genes or regulatory sequences.

Nanopore long reads and Flye assembly are a low-cost, rapid, method of producing quality genomes. The cost and speed at which these analyses can be done may allow an expansion of insect genome sequencing, allowing us to better probe the diversity of genomes at large scale in this most successful group of animals.

We hope the ease of this method, in addition to the sharing of our scripts, will the genomics community to rapidly update assembly methods in real time to take full advantage of updated technologies.

## Data availability

The genomes described here are available at NCBI with the following accession numbers.

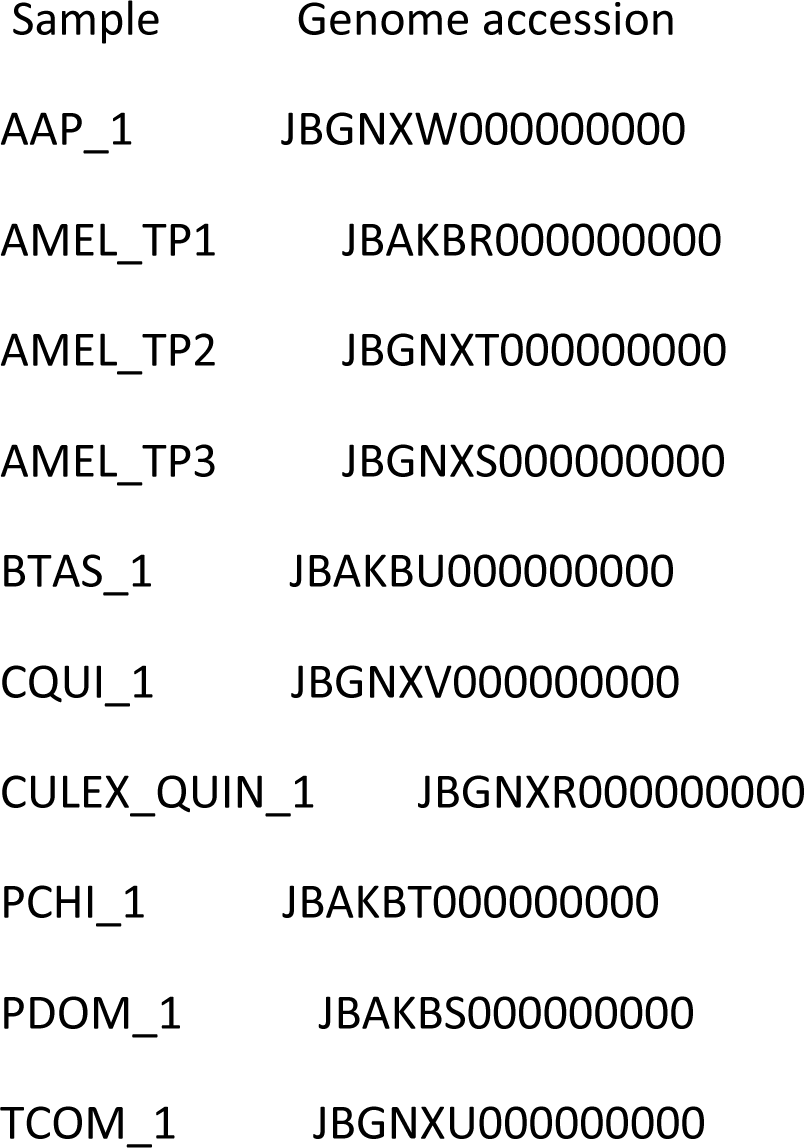

| Sample | Genome accession |
| --- | --- |
| AAP_1 | JBGNXW000000000 |
| AMEL_TP1 | JBAKBR000000000 |
| AMEL_TP2 | JBG NXT000000000 |
| AMEL_TP3 | JBG NXS000000000 |
| BTAS_1 | JBAKBU000000000 |
| CQUI_1 | JBG NXV000000000 |
| CULEX_QUIN_1 | JBG NXR000000000 |
| PCHI_1 | JBAKBT000000000 |
| PDOM_1 | JBAKBS000000000 |
| TCOM_1 | JBG NXU000000000 |

## Acknowledgements

*Celatoblatta quinquemaculata* were collected from the Rock and Pillar Range by Dr Craig Marshall. *Anatalanta aptera* was kindly provided by David Renault, University of Rennes. This work was funded by Genomics Aotearoa.

## Author Contributions

**Conceptualization: P.K.D**., **T.H.; Methodology: J.G, T.H. S.N.I. P.K.D**., **P.K**., **K.M**., **J.G.; Validation: J.G**., **P.K.D.; Formal analysis: J.G**., **J.G**., **S.N.I**., **P.K.D.; Resources: P.K.D**., **C.J.S**., **D.W**., **A.H**., **G.P**., **J.K. ; Data curation: J.G**., **J.G. ; Writing - original draft: J.G**..; P.K.D. Visualization: J.G. Supervision: P.K.D.; Project administration: P.K.D.; Funding acquisition: P.K.D.

## Supplemental Figure and Tables

**Supplemental Figure 1:**
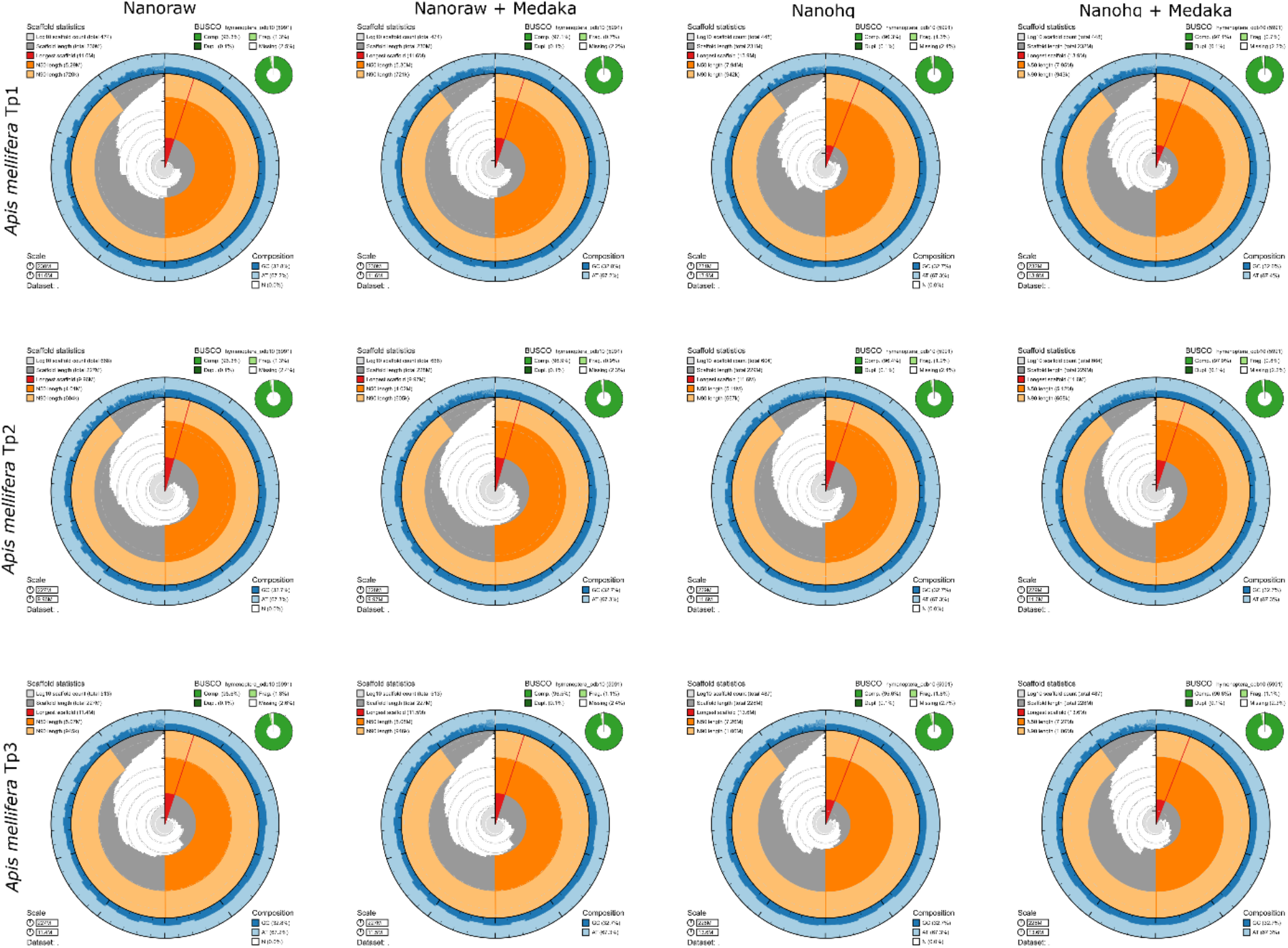
BlobToolKit Snail plot of honeybee drone assemblies, from three drones TP1, TP2 and TP3, generated using different approaches as labelled.

**Supplemental Figure 2:**
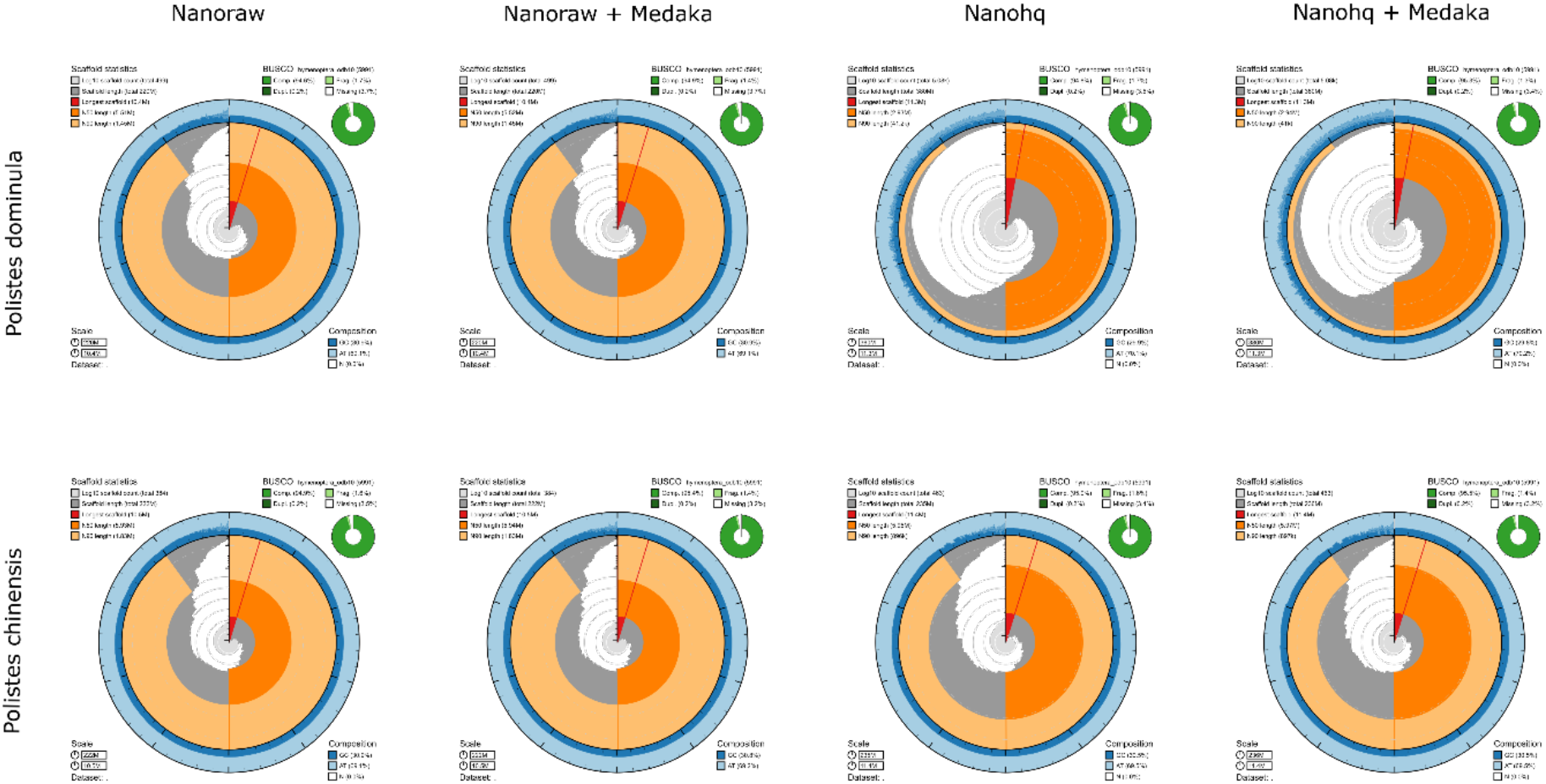
BlobToolKit Snail plot of *Polistes* drone assemblies, from *P. dominula* and *P. chinensis*, generated using different approaches as labelled. Note the nanohq approachs on *P*.*s chinensis* produced a doubled-sized genome.

**Supplemental Figure 3:**
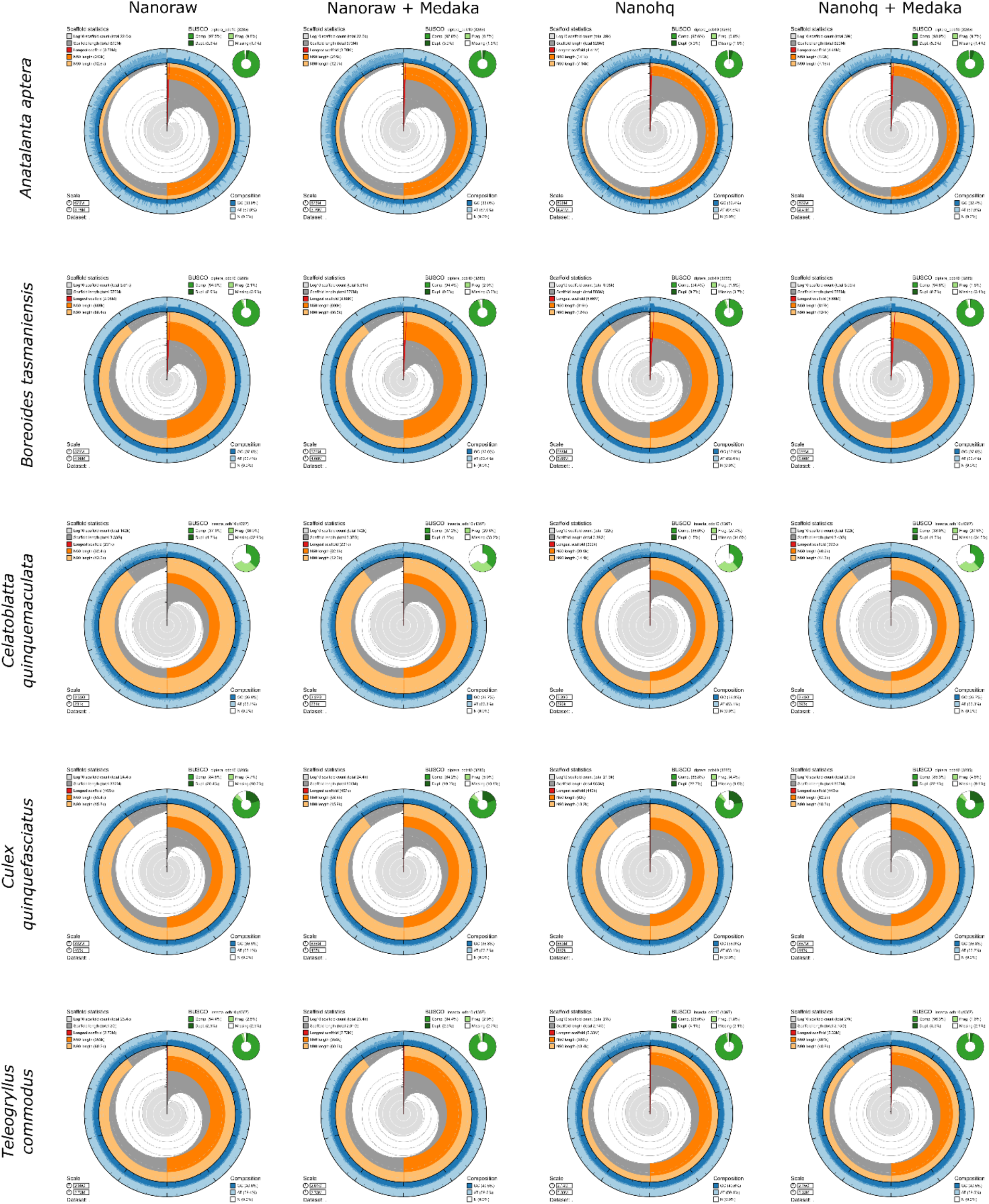
BlobToolKit Snail plot of assemblies of diploid insects generated using different approaches as labelled.

**Supplemental Table 1:** Honeybee BUSCO scores.

| Name | BUSCO Lineage | BUSCO Complete | BUSCO Duplicated | BUSCO Fragmented | BUSCO Missing | BUSCO lineage Total |
| --- | --- | --- | --- | --- | --- | --- |
| TP1 hq | hymenoptera_odb10 | 96.32782507 | 0.06677 | 1.25187782 | 2.420297112 | 5991 |
| TP1 hq medaka | hymenoptera_odb10 | 97.12902687 | 0.05008 | 0.70105158 | 2.169921549 | 5991 |
| TP1 raw | hymenoptera_odb10 | 96.26105825 | 0.06677 | 1.28526123 | 2.453680521 | 5991 |
| TP1 raw medaka | hymenoptera_odb10 | 97.09564347 | 0.06677 | 0.68435987 | 2.219996662 | 5991 |
| TP2 hq | hymenoptera_odb10 | 96.36120848 | 0.06677 | 1.23518611 | 2.403605408 | 5991 |
| TP2 hq medaka | hymenoptera_odb10 | 96.97880154 | 0.06677 | 0.7845101 | 2.236688366 | 5991 |
| TP2 raw | hymenoptera_odb10 | 96.26105825 | 0.08346 | 1.30195293 | 2.436988817 | 5991 |
| TP2 raw medaka | hymenoptera_odb10 | 96.87865131 | 0.06677 | 0.85127692 | 2.270071774 | 5991 |
| TP3 hq | hymenoptera_odb10 | 95.59339009 | 0.06677 | 1.75262894 | 2.653980971 | 5991 |
| TP3 hq medaka | hymenoptera_odb10 | 96.61158404 | 0.05008 | 1.06826907 | 2.320146887 | 5991 |
| TP3 raw | hymenoptera_odb10 | 95.61008179 | 0.08346 | 1.76932065 | 2.620597563 | 5991 |
| TP1 hq | hymenoptera_odb10 | 96.32782507 | 0.06677 | 1.25187782 | 2.420297112 | 5991 |
| **Amel_HAv3.1** | hymenoptera_odb10 | 97.42947755 | 0.06677 | 0.33383408 | 2.236688366 | 5991 |

**Supplemental Table 2:** *Polistes* assembly BUSCO scores.

| Name | BUSCO Lineage | BUSCO Complete | BUSCO Duplicated | BUSCO Fragmented | BUSCO Missing | BUSCO lineage Total |
| --- | --- | --- | --- | --- | --- | --- |
| Pchi hq | hymenoptera_odb10 | 94.77549658 | 0.2003 | 1.70255383 | 3.521949591 | 5991 |
| Pchi hq medaka | hymenoptera_odb10 | 95.259556 | 0.23368 | 1.31864463 | 3.421799366 | 5991 |
| Pchi raw | hymenoptera_odb10 | 94.57519613 | 0.2003 | 1.71924553 | 3.705558338 | 5991 |
| Pchi raw medaka | hymenoptera_odb10 | 94.94241362 | 0.23368 | 1.36871975 | 3.688866633 | 5991 |
| Pdom hq | hymenoptera_odb10 | 94.99248873 | 0.21699 | 1.60240361 | 3.405107661 | 5991 |
| Pdom hq medaka | hymenoptera_odb10 | 95.45985645 | 0.21699 | 1.35202804 | 3.188115507 | 5991 |
| Pdom raw | hymenoptera_odb10 | 94.94241362 | 0.23368 | 1.55232849 | 3.505257887 | 5991 |
| **GCF_001465965.1**<br>**Pdom Reference**<br>**Genome** | hymenoptera_odb10 | 95.61008179 | 0.25038 | 0.98481055 | 3.405107661 | 5991 |

**Supplemental Table 3:** Diploid insect assembly BUSCO scores.

| Name | Lineage | Complete | Duplicated | Fragmented | Missing | Total |
| --- | --- | --- | --- | --- | --- | --- |
| *Antalanta* hq | diptera_odb10 | 97.62557 | 5.266362 | 0.791476 | 1.582952816 | 3285 |
| *Antalanta* hq medaka | diptera_odb10 | 97.96043 | 5.296804 | 0.669711 | 1.369863014 | 3285 |
| *Antalanta* raw | diptera_odb10 | 97.50381 | 3.866058 | 0.821918 | 1.674277017 | 3285 |
| *Antalanta* raw medaka | diptera_odb10 | 97.80822 | 3.896499 | 0.669711 | 1.522070015 | 3285 |
| *Boreoides* hq | diptera_odb10 | 94.42922 | 0.700152 | 1.887367 | 3.683409437 | 3285 |
| *Boreoides* hq medaka | diptera_odb10 | 94.79452 | 0.730594 | 1.796043 | 3.409436834 | 3285 |
| *Boreoides* raw | diptera_odb10 | 94.00304 | 0.608828 | 2.100457 | 3.896499239 | 3285 |
| *Boreoides* raw medaka | diptera_odb10 | 94.39878 | 0.608828 | 1.978691 | 3.622526636 | 3285 |
| *Teleogryllus* hq | insecta_odb10 | 96.04974 | 4.096562 | 1.828822 | 2.121433797 | 1367 |
| *Teleogryllus* hq medaka | insecta_odb10 | 96.2692 | 3.511339 | 1.609364 | 2.121433797 | 1367 |
| *Teleogryllus* raw | insecta_odb10 | 94.36723 | 2.926116 | 2.77981 | 2.852962692 | 1367 |
| *Teleogryllus* raw medaka | insecta_odb10 | 94.36723 | 2.487198 | 2.926116 | 2.706656913 | 1367 |
| *Culex* hq | diptera_odb10 | 85.81431 | 22.70928 | 4.383562 | 9.802130898 | 3285 |
| *Culex* hq medaka | diptera_odb10 | 85.32725 | 22.10046 | 4.809741 | 9.863013699 | 3285 |
| *Culex* raw | diptera_odb10 | 84.59665 | 20.42618 | 4.718417 | 10.68493151 | 3285 |
| *Culex* raw medaka | diptera_odb10 | 84.17047 | 19.33029 | 5.022831 | 10.80669711 | 3285 |
| *Celatoblatta* hq | insecta_odb10 | 38.0395 | 1.463058 | 27.35918 | 34.60131675 | 1367 |
| *Celatoblatta* hq medaka | insecta_odb10 | 37.96635 | 1.536211 | 27.57864 | 34.45501097 | 1367 |
| *Celatoblatta* raw | insecta_odb10 | 37.08851 | 1.170446 | 29.99268 | 32.91880029 | 1367 |
| *Celatoblatta* raw medaka | insecta_odb10 | 37.16167 | 1.316752 | 29.62692 | 33.21141185 | 1367 |

